# *Hanseniaspora* yeasts from agave fermentations provide insight into intraspecific gene loss

**DOI:** 10.1101/2025.10.24.684411

**Authors:** Luis F. García-Ortega, Ángela M. García-Acero, Manuel R. Kirchmayr, Lucia Morales, Alexander DeLuna, Luis Delaye, Eugenio Mancera

## Abstract

*Hanseniaspora* species stand out among yeasts for having the smallest genomes, marked by extensive gene family contractions. Consequently, they have been proposed as model organisms for studying the evolution of free-living cells lacking genes otherwise considered essential. Here, we show that *Hanseniaspora* yeasts are prevalent in agave fermentations used to produce traditional spirits across Mexico. We sequenced the genomes of 15 strains, unambiguously identifying them as *H. lachancei, H. pseudoguilliermondii, H. guilliermondii*, and *H. opuntiae*. Comparative genomic analyses revealed dynamic shifts in gene family sizes and compositions, suggesting ongoing gene loss within the fast-evolving branch of this genus. Notably, gene losses varied across functional categories, even among isolates of the same species. Growth assays across diverse stress conditions and carbon sources indicated that these genomic disparities do not directly translate into phenotypic differences. Together, our findings suggest that differential gene loss within species is an active evolutionary process shaping *Hanseniaspora* genomes, highlighting agave-fermentation populations as valuable models for studying genome reduction in a natural context.

## INTRODUCTION

Yeasts from the genus *Hanseniaspora* (anamorph *Kloeckera*) are a group of ascomycetes fungi known for their apiculate cellular shape and their role in wine fermentation. Although their fermentation capacity is weaker than that of *Saccharomyces* yeasts, these microorganisms can thrive in wine fermentations, especially in the initial phases. These fungi produce a variety of volatile compounds, such as esters, aldehydes, terpenes and higher alcohols, which contribute fruity, floral, or spicy notes to fermented beverages (González-Robles et al., 2015). Additionally, some studies suggest that *Hanseniaspora* yeasts may exert a protective effect against spoilage organisms, potentially contributing to the stability and longevity of wine and other fermented products (Xu et al., 2006; Gomomo et al., 2022). These microorganisms are cosmopolitan and are commonly found on plant surfaces and insects, which may act as reservoirs for open fermentations (Kurtzman et al., 2011).

Apart from their roles in open fermentations, *Hanseniaspora* yeasts have drawn attention due to their rapid evolution and high rates of gene loss. They have the smallest genomes among yeasts (∼4,000 compared to ∼6,000 genes) and lack genes from pathways that are central for cellular physiology such as cell-cycle control and DNA repair (Steenwyk et al., 2019; Haase et al., 2024). Based on their evolutionary rate, the 24 known *Hanseniaspora* species (Ryan et al., 2024) have been categorized into two distinct groups, a fast-evolving and a slow-evolving lineage (FEL and SEL, respectively) (Steenwyk et al., 2019). Taking advantage of these unusual genomic characteristics, *Hanseniaspora* yeasts have been employed as a model to explore the long-term effects of high mutation rates on eukaryotic genomes (Steenwyk et al., 2019; Saubin et al., 2020; Haase et al., 2024; Onetto et al. 2025).

Recent surveys of the microbial communities that are involved in the open fermentations used to produce traditional agave spirits in Mexico have shown that *Hanseniaspora* are common in such environments. These fermentations are dynamic ecosystems teeming with a diverse community of microorganisms, mostly yeasts and bacteria (Jara-Servin et al., 2025; Nolasco-Cancino et al., 2018; Torres-Velázquez et al., 2022; Colón-González et al., 2025). The microorganisms play a pivotal role in the production process, converting the sugars in cooked agave to ethanol and other compounds that contribute to the distinctive flavor and aroma of agave spirits (Lappe-Oliveras et al., 2008). *Hanseniaspora* yeasts are part of the core fungal species of this environment (Lachance, 1995; Gallegos-Casillas et al., 2024; Jara-Servin et al., 2025), with *H. lachancei* reported so far only from this setting (Lachance, 1995). It is not yet clear whether *H. lachancei* is endemic to the geographic region and the agave fermentation environment since it is a relatively understudied species with limited number of strains reported and available in culture collections. Moreover, the identification of *H. lachancei* is challenging because it belongs to a species complex with three close relatives (*H. guilliermondii, H. pseudoguilliermondii* and *H. opuntiae*) (Cadez et al. 2003), from which only *H. guilliermondii* has been also reported in agave fermentations (Kirchmayr et al. 2017; Gallegos-Casillas et al., 2024; Jara-Servin et al., 2025).

In this study, we characterized the genomic and phenotypic diversity of *Hanseniaspora* isolates from open agave fermentations. Analysis of the genomic sequences showed that the loss of genes is an ongoing process, as we identified differences in gene content between strains of the same species. However, in general, these differences did not correlate with the phenotypic profile of the strains in various carbon sources and stress conditions. Our results raise the question of whether environmental factors are driving the patterns of gene loss observed in these species, highlighting agave fermentation isolates as a powerful model for understanding genome contractions within species.

## MATERIALS AND METHODS

### Strain identification by ITS sequencing

We selected a panel of 15 strains from the YMX-1.0 culture collection (Gallegos-Casillas et al., 2024) identified by MALDI-TOF as *Hanseniaspora*. Isolates were restreaked twice to obtain axenic (pure) cultures, and genomic DNA was extracted using sodium hydroxide. The ITS/5.8 rDNA region was amplified and sequenced with the universal fungal primers ITS-1 and ITS-4, following the method described by (Sylvester et al. 2015). Sequences were filtered with a 0.01 trimming threshold and assembled using a modified Mott (M1) cut algorithm, as implemented in the *sangeranalyseR* (Chao et al. 2021) R package. The assembled sequences were then compared to annotated yeast sequences in the GenBank database using BLAST (Altschul et al. 1990).

### Genome Sequencing, *de novo* assembly and annotation

Genomic DNA was purified using the MasterPure DNA purification kit as recommended by the manufacturer, and it was sequenced at BGI (Shenzhen, China) using the DNBSeq platform using 150bp paired-end reads. Raw reads were quality assessed using fastp v0.20.0 (Chen et al., 2018) to remove adapter sequences and low-quality reads. *de novo* assembly of the filtered reads was performed using SPAdes v3.12.0 (Bankevich et al. 2012) with the BayesHammer module for error correction and iterative k-mer lengths (21, 33, 55, and 77□bp). After assembly, iterative scaffolding with RagTag (v2.1.0) (Alonge et al. 2022) and manual curation was performed using the *H. opuntiae* (strain AWRI3578) genome as reference. The final assemblies were annotated using the MAKER pipeline (v3.01.3) (Cantarel et al. 2008). A first round of MAKER was computed to construct gene models directly from both aligned transcript sequences and reference proteins. Then, two additional rounds of annotation using MAKER with Snap v2013-02-16 (Korf 2004), Augustus v3.2.1 (Stanke et al., 2006) and evidence build (proteins and transcripts), were performed to create an *ab initio* evidence-driven gene build. For homology-based predictions we used a custom deduplicated data set of peptide and transcript sequences from *H. opuntiae* strain AWRI3578, *H. osmophila* strain AWRI3579 and *H. meyeri* strain APC 12.1 (Table S2). For *ab initio* gene predictions, we used a gene set with the best gene models based on Annotation Edit Distance (AED) scores < 0.25 and a minimal amino acid length of 50. Augustus was trained with BUSCO (Simão et al., 2015) using the Saccharomycetes odb10 dataset. Protein and gene names were assigned to the gene predictions using a BLASTp v.2.7.1 (Altschul et al. 1990) search of predicted protein sequences against the UniProt/Swiss-Prot database with an E-value < 10^-6^.

### Pseudogene identification

To identify pseudogenes and verify that apparent gene absences were not artifacts of false stop codons introduced during sequencing or assembly, we used Pseudopipe (Zhang et al., 2006) on the genome drafts with default parameters. The protein sequences of the deduplicated data set described in the annotation process were employed as queries. Putative pseudogenes were filtered by excluding those that overlapped with gene annotations, transposon elements, or sequences shorter than 150 bp.

### GO enrichment analysis

To identify enriched functional categories among absent, present, and pseudogenized genes, Gene Ontology (GO) for each predicted protein sequence in the *Hanseniaspora* genomes were obtained using InterProScan v.5.35-74.0 (Jones et al., 2014) with the Pfam library. This information was used to create a customized GO database for each species using the AnnotationForge R package (Carlson and Herves, 2023). GO enrichment analysis was performed with Fisher’s exact test as implemented in the topGo package (Alexa and Rahnenfuhrer, 2024), using the classic algorithm, and a significance cutoff *p*-value of < 0.01. Reduction and visualization of GO Biological Process terms were performed using the R package rrvgo v4.5 (Sayols, 2023). Term similarity was computed with calculateSimMatrix based on the *S. cerevisiae* annotation database (org.Sc.sgd.db). Representative terms were selected using reduceSimMatrix with a similarity threshold of 0.7. The resulting terms were visualized via UMAP, where point size indicates the number of genes associated with each GO term.

### Phylogenetic analysis

Phylogenomic analysis was performed using the strains sequenced and 16 representative *Hanseniaspora* species with genomic sequences available in the NCBI database (Table S1 and S2). To prevent a bias by genome annotation, all genomes were annotated with MAKER as described for the strains here sequenced. The species tree and gene orthogroups were inferred using the STRIDE and STAG algorithms implemented in OrthoFinder as previously described (Emms and Kelly, 2019). *Saccharomyces cerevisiae* S288C was used as the outgroup.

### Phenotyping assays

Fifteen strains isolated from agave fermentations and 17 strains from the ARS culture collection (NRRL) covering most known species for the *Hanseniaspora* genus (Table S3) were subjected to high throughput phenotyping by micro-cultivation in different growth conditions (Table S4), as previously described (Espinosa-Cantú et al., 2018). *S. cerevisiae* S288c was used as a reference strain. Strains were inoculated in 100 μl of Yeast extract-Peptone-Dextrose medium (YPD: 1% yeast extract, 2% peptone, and 2% dextrose) and incubated for 48 h at 30°C. Growth was monitored in an automated robotic system (Tecan Freedom EVO200) that integrates a plate carrousel (Liconic STX110), a plate reader (Tecan Infinite M1000), an orbital plate shaker, and a robotic manipulator arm. The equipment was maintained in an environmental room at constant temperature (30°C). Absorbance at 600 nm (OD600) was measured every 60 min after vigorous shaking, and growth kinetics were followed for 48 hours. The culture medium without inoculum was used as a blank control. To test variation within plates, four of the strains (YMX002864, YMX005643, Y-27514 and S288c) were grown in triplicate in each of the conditions evaluated.

## RESULTS

### *Hanseniaspora* yeasts are prevalent in agave fermentations

Metabarcoding efforts have shown that *Hanseniaspora* yeasts are part of the core fungal microbiome of agave fermentations, being present in at least 90% of the distilleries studied and in all of the seven agave spirit producing regions in Mexico (Jara-Servin et al., 2025). In terms of isolates, a total of 160 *Hanseniaspora* strains were initially identified using MALDI-TOF biotyping as part of the YMX□1.0 culture collection of yeasts from traditional agave fermentations (Gallegos-Casillas et al., 2024). These strains originated from 15 distilleries located across four states within five of the seven regions where agave spirits are produced in Mexico: Balsas basin, Northwest, South central, West I, and West II (Figure 1-A). No *Hanseniaspora* strains were isolated in the Northeast or Central Highlands regions, even when metabarcoding had shown that these yeasts are present (Jara-Servin et al., 2025). According to the identification by MALDI-TOF, these 160 isolates belonged to four species: *H. lachancei* (47.5%), *H. valbyensis* (29.4%), *H. opuntiae* (22.5%) and *H. vineae* (0.6%). The South-central region was the only region in which these four species were isolated, with *H. lachancei* being the most abundant. *H. opuntiae* was the only species isolated in the five regions in which *Hanseniaspora* strains were recovered, with the Northwest being the region where it is relatively most abundant.

**Figure 1.**
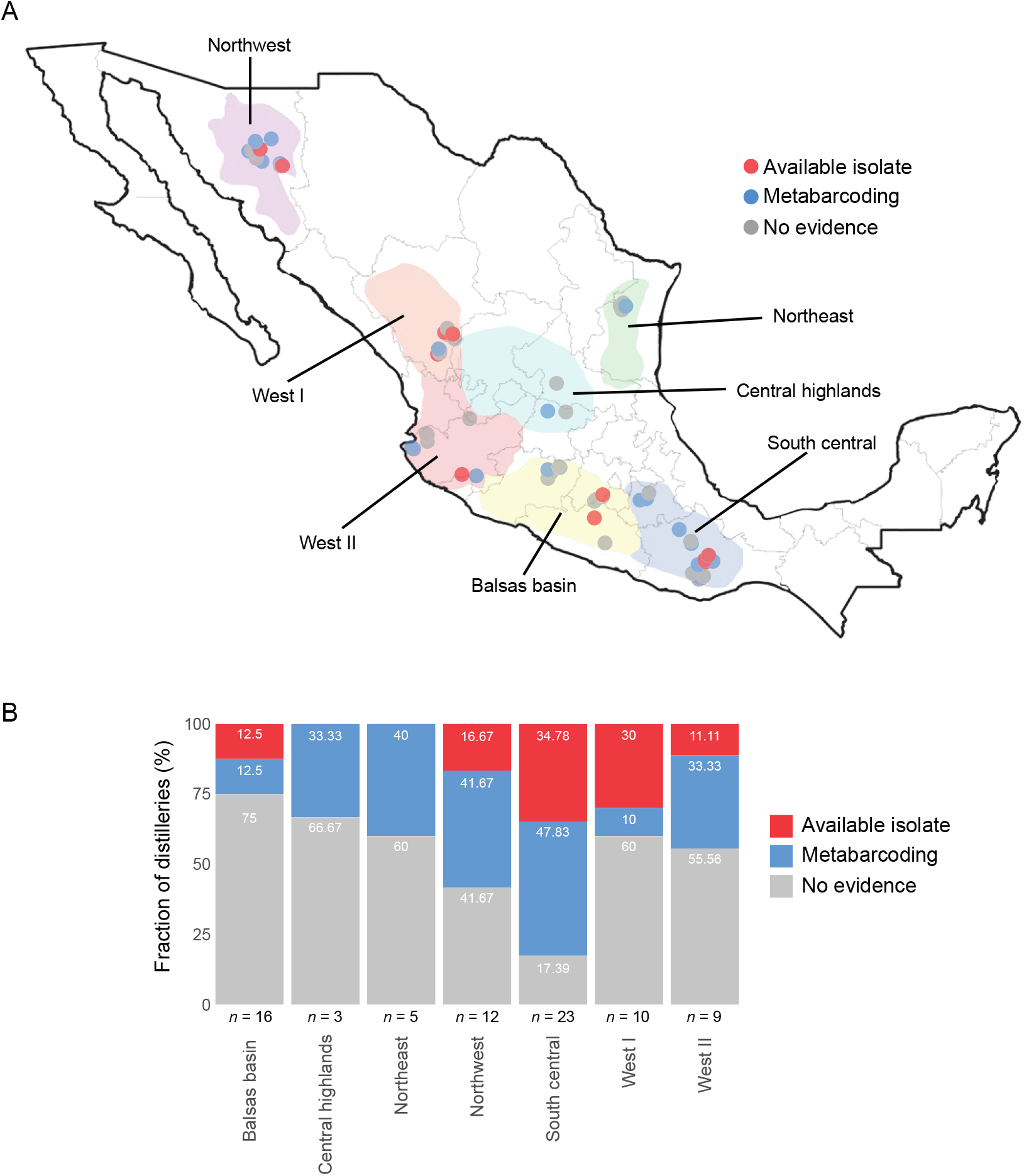
Distribution of *Hanseniaspora* isolates across agave fermentation in Mexico. A) *Hanseniaspora* species were isolated from five of the agave spirit producing regions (red dots) and species from the genus were detected by metabarcoding in six of the seven regions (blue dots). Grey dots indicate sampled distilleries, with no evidence of *Hanseniaspora* presence by either detection method. The map is taken from Gallegos-Casillas et al., (2024), showing specifically the distribution of *Hanseniaspora* species including data from Gallegos-Casillas et al., (2024) and Jara-Servin et al. (2025). B) Stacked bar plot illustrates the fraction (%) of distilleries in each region where there is evidence of *Hanseniaspora* presence by metabarcoding (blue) or strain isolation (red). The number of distilleries sampled in each region (*N)* is indicated for each bar.

To verify the strain identity of the *Hanseniaspora* isolates from agave fermentations, we performed ITS sequencing of 15 of these strains. The results showed discrepancies with the species assigned by MALDI-TOF biotyping (Table S1), although both methods were highly accurate at the genus level. By ITS sequencing, the *Hanseniaspora* species identified were *H. lachancei, H. opuntiae, H. guilliermondii* and *H. osmophila*. Metabarcoding analysis of agave fermentations also based on sequencing of the ITS region had previously found *H. valbyensis, H. uvarum, H. osmophila* and a species only identified at the genus level (Jara-Servin et al., 2025). Yet, species assignment based on whole genome sequencing, as detailed below, showed inaccuracies in the identification by both MALDI-TOF biotyping and ITS sequencing (Table S1). As expected, ITS sequencing was more accurate, but failed to identify *H. pseudoguilliermondii*, one of the four members of the *H. lachancei - H. guilliermondii - H. pseudoguilliermondii - H. opuntiae* complex. Overall, besides underscoring the challenges of identifying members of this species complex, our results showed that agave fermentations in Mexico represent a common habitat for *Hanseniaspora* yeasts.

### Genomic diversity of *Hanseniaspora* isolates from agave fermentations

To assess genomic diversity within *Hanseniaspora* isolates from agave fermentations, we selected 15 representative strains for whole-genome sequencing, 13 of the strains that had been identified by ITS sequencing (Table S1), together with two additional strains previously isolated (YMX005643 and YMX005644). We mainly focused on isolates of *H. lachancei*, the only specie so far found to be endemic to agave fermentations, but also included representatives of the three other members of the species complex. In total we sequenced nine *H. lachancei* (five from South Central, two from West I and two from West II), three *H. pseudoguilliermondii* (two from South Central and one from West I), two *H. guilliermondii* (one from South Central and one from the Balsas Basin), and one *H. opuntiae* from West II. Assembly and scaffolding of the 15 genomes yielded between 28 and 1,116 scaffolds per genome (Materials and Methods). The total genome sizes of the assemblies ranged from 8,711,511 to 9,069,783 bp, with an average size of 8,817,845 bp (Table S5). These values are consistent with the reference genome sizes reported for the species within the *H. lachancei* complex: 9.14 Mb (*H. guilliermondii*), 8.90 Mb (*H. lachancei*), 8.79 Mb (*H. pseudoguilliermondii*), and 8.83 Mb (*H. opuntiae*). The average number of predicted genes in the 15 sequenced strains was 4,162, with an average coding sequence length of 1,500 bp and 1.17 exons per gene. This also aligns with the average number of genes reported for the *Hanseniaspora* genus, which is 4,708 ± 634 genes (Rueda-Mejia et al., 2023).

To evaluate genome assembly completeness, we compared our assemblies to a set of 2,137 Benchmarking Universal Single-Copy Orthologs (BUSCOs) (Simão et al., 2015). This analysis revealed an average coverage of 58.9% complete BUSCOs in our *Hanseniaspora* assemblies. This value increased to 79.8% following gene annotation. The low level of BUSCO coverage that we observed was not unexpected, given the genomic contractions known to have occurred in these yeasts. For example, if we calculate BUSCO coverage of strains previously sequenced by long-reads and whose genomes are assembled at the chromosome level we estimated around 79% coverage (Rueda-Mejia, et al., 2023; Cadez et al., 2021). Additionally, we identified between 143 and 306 kb of repetitive sequences within the *Hanseniaspora* genomes that we sequenced, representing between 1.6% and 3.4% of each assembly.

Phylogenetic analysis of the 15 sequenced agave fermentation isolates along with representative *Hanseniaspora* genomes, utilizing 5,429 gene families, confirmed the expected taxonomic placement of the strains (Figure 2). This phylogeny was consistent with previously established relationships for the *Hanseniaspora* species (Steenwyk et al., 2019). We identified the slow- and fast-evolving lineages (SEL and FEL, respectively) and, importantly, confirmed the close phylogenetic relationships within the *H. opuntiae* - *H. lachancei* - *H. guilliermondii* - *H. pseudoguilliermondii* species complex. The close relationship observed between *H. opuntiae* and *H. pseudoguilliermondii* is also supported by previous research (Saubin et al., 2020; Onetto et al., 2025).

**Figure 2.**
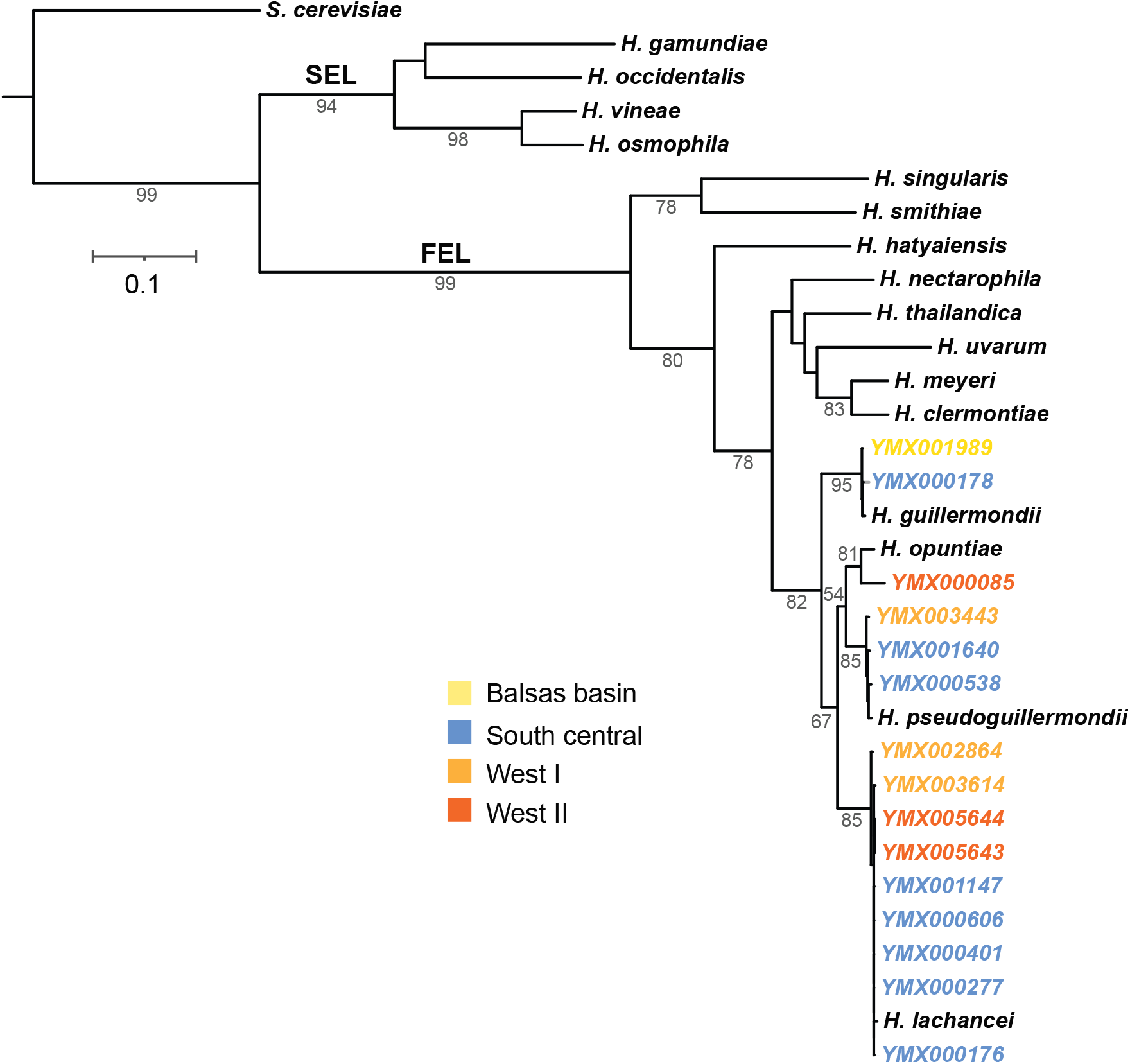
Phylogenetic relationships among *Hanseniaspora* isolates from agave fermentations. The phylogenetic tree was inferred from 5,429 orthogroups using the STRIDE and STAG algorithms as implemented in the OrthoFinder program. Support values represent the proportion of times that the bipartition is observed in each of the individual species tree estimates. Only values exceeding 50% are shown. Branch lengths indicate the average number of substitutions per site across the sampled gene families. *S. cerevisiae* served as the outgroup. The tree highlights the FEL and SEL lineages, as well as the geographic origin of each isolate with different label colors.

### Intraspecies gene-loss variability within *Hanseniaspora* isolates

Evolutionary gene loss is a hallmark of *Hanseniaspora* species (Steenwyk et al., 2019), a trend confirmed by our analysis of isolates from agave fermentations. However, it remains unclear whether this observed losses stem from a single event in the FEL ancestor or represent an ongoing event across the phylogeny. The abundance of *Hanseniaspora* isolates in our collection enabled us to interrogate this phenomenon in a natural fermentation context.

By comparing protein families, we identified 5,298 orthogroups (OGs) among the 31 sequenced and reference *Hanseniaspora* genomes (excluding *S. cerevisiae*), which constitute the genus’s pan-genome. Of these, 2,111 OGs were conserved with at least one gene present in all 31 genomes, forming the core-genome. Furthermore, the pan-genome of the *H. opuntiae* - *H. lachancei* - *H. guilliermondii* - *H. pseudoguilliermondii* species complex comprises 4,189 OGs, representing 79.18% of the genus pan-genome. Additionally, for 3,854 (92%) of these OGs, at least one gene copy was present across all isolates. This suggests that the remaining 335 OGs exhibited a scattered distribution among the genomes of the species complex, including the isolates from agave fermentations.

Clustering analysis revealed substantial intraspecific variability in gene loss patterns within some *Hanseniaspora* species (Figure 3). While all genomes of *H. guilliermondii* and *H. pseudoguilliermondii* show similar patterns and therefore cluster together according to gene loss, isolates from *H. opuntiae* were situated at two distant positions along the dendrogram. Similarly, *H. lachancei* isolates YMX005643 and YMX005644 exhibit higher degree of dissimilarity to all other *Hanseniaspora* species, including their conspecific isolates. The observed within-species diversity in gene losses suggests that *Hanseniaspora* species continue to evolve through gene loss.

**Figure 3.**
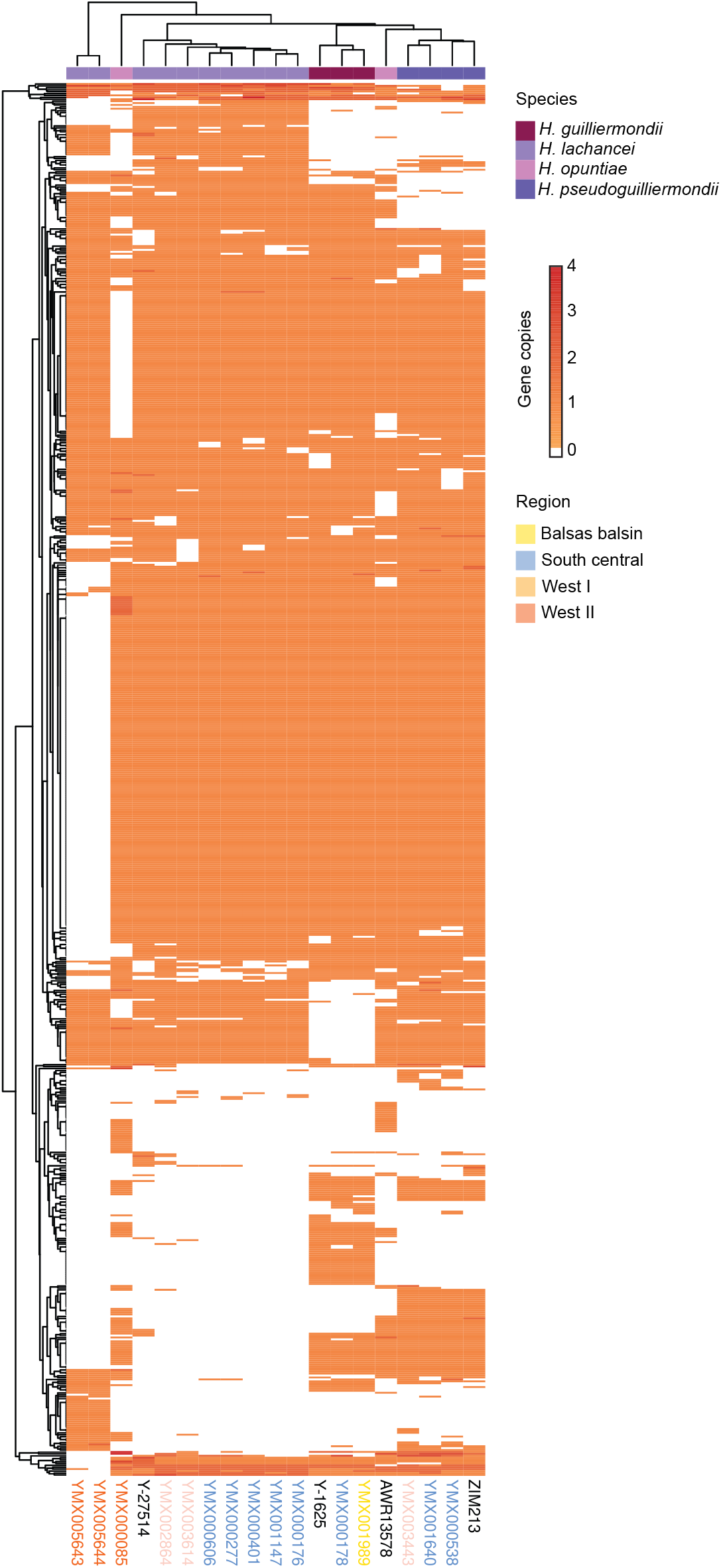
Gene loss varies among *Hanseniaspora* isolates from agave fermentations. Heatmap showing unsupervised clustering based on genome-wide gene presence (orange) and absence (white) across the sequenced *Hanseniaspora* isolates. Previously published genomes were included as references. The colored track above the heatmap indicates the species of each strain, while the color of the strain names (below) denotes the geographic region of origin. The labels of the reference strains are shown in black.

Gene Ontology (GO) enrichment analysis of gene losses further showed that they are not uniformly distributed among isolates of the same species (Figure 4). For instance, neither *H. guilliermondii* nor *H. pseudoguilliermondii* clustered together in the dendrogram, indicating different patterns of enrichment. Differences were also evident in the extent of gene loss across categories; for example, isolates YMX000085 and AWR13578 of *H. opuntiae* displayed markedly different patterns. These specific isolates showed a differential depletion of genes involved in cell cycle and mitotic cell cycle, cellular response to stimulus, ribonucleoprotein complex, DNA repair and DNA damage response, cellular response to stress, and macromolecule metabolic process. Similarly, *H. lachancei* isolates showed within species variability in gene losses related to several different categories. This pattern reveals that *Hanseniaspora* lineages exhibit varying pathways within species where gene loss has occurred.

**Figure 4.**
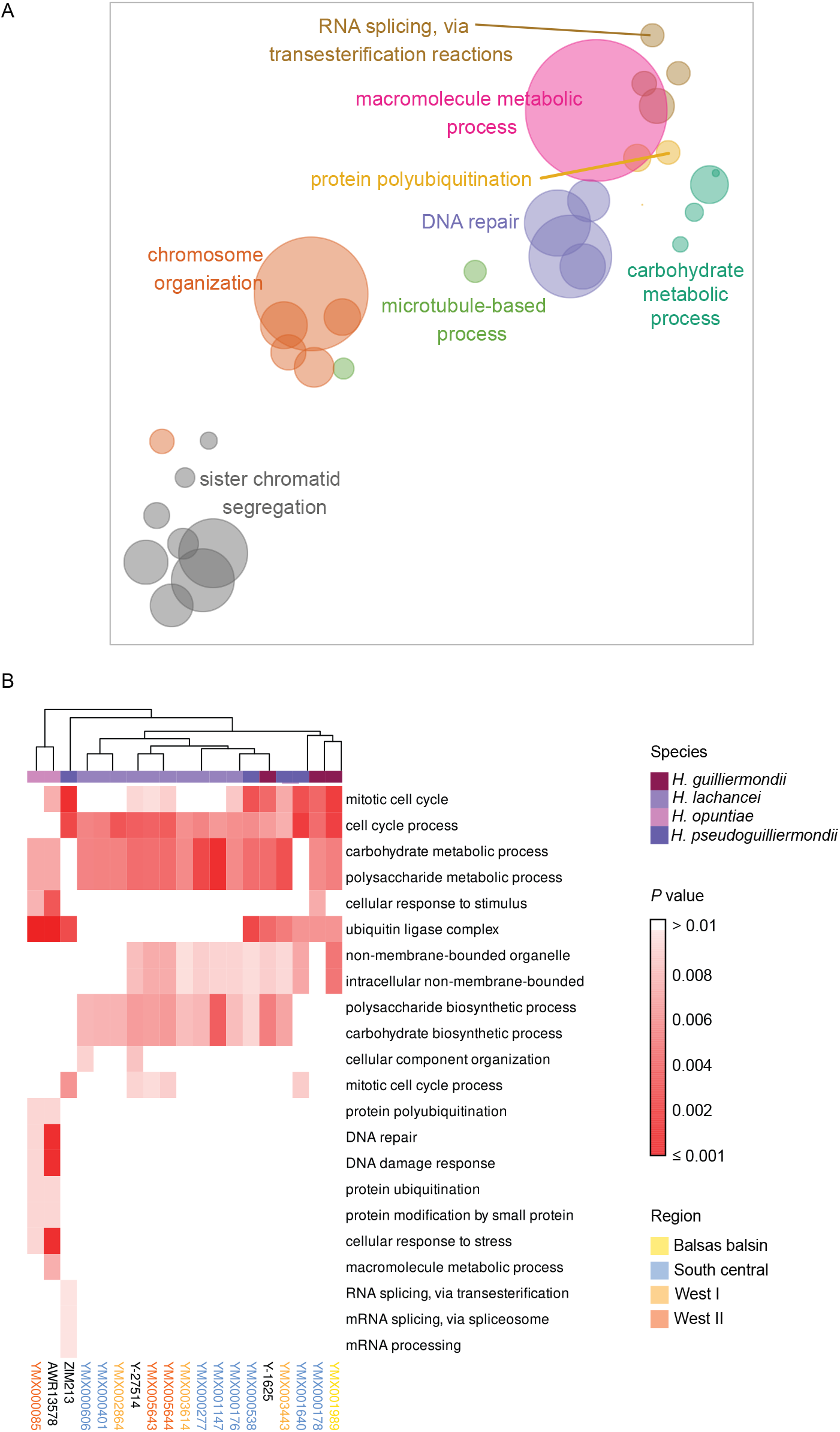
Enriched GO categories associated with gene losses in *Hanseniaspora* isolates. GO enrichment analysis of biological processes was performed using Fisher’s exact test with a significance threshold of *p* < 0.01. A) Dispersion plot showing GO biological processes enriched among missing genes in the sequenced *Hanseniaspora* isolates and reference genomes, after reducing term redundancy with rrvgo. Each circle represents a cluster of redundant GO terms, colored by broader superclusters of related functions. Distances between circles reflect term similarity, with axes corresponding to the first two UMAP components of the dissimilarity matrix. B) Heatmap showing the GO terms that are variable among isolates. Red to withe colors indicate the *P* value of the enrichment as indicated in the legend. Categories that were not statistically significant are shown in white. The colored track above the heatmap indicates the species of each strain, while the color of the strain names (below) denotes the geographic region of origin. The labels of the reference strains are shown in black.

### Phenotyping analysis of *Hanseniaspora* isolates from agave fermentations

To investigate how specific gene loss patterns might influence phenotypic traits, we assessed the fitness of each *Hanseniaspora* strain under diverse stress conditions and across different carbon and nitrogen sources (Table S4). In addition to the strains isolated from agave fermentations, we included representatives of most other *Hanseniaspora* species (Table S3). Fitness was measured as the ratio of the growth rate under each condition to the growth rate in rich medium as a reference condition. Hierarchical clustering of the measured growth rates identified two primary clusters in terms of the growth conditions. The first cluster comprised conditions with generally low fitness, including most tested carbon sources (excluding fructose) and salt stress conditions (except CaCl_2_ at 200 mM). The second cluster was defined by growth under nitrogen sources such as urea, tyrosine, methionine, and phenylalanine, and stress conditions like 6-azauracil in which strains from *H. lachancei* and *H. pseudoguilliermondii* exhibited higher fitness (Figure 5).

**Figure 5.**
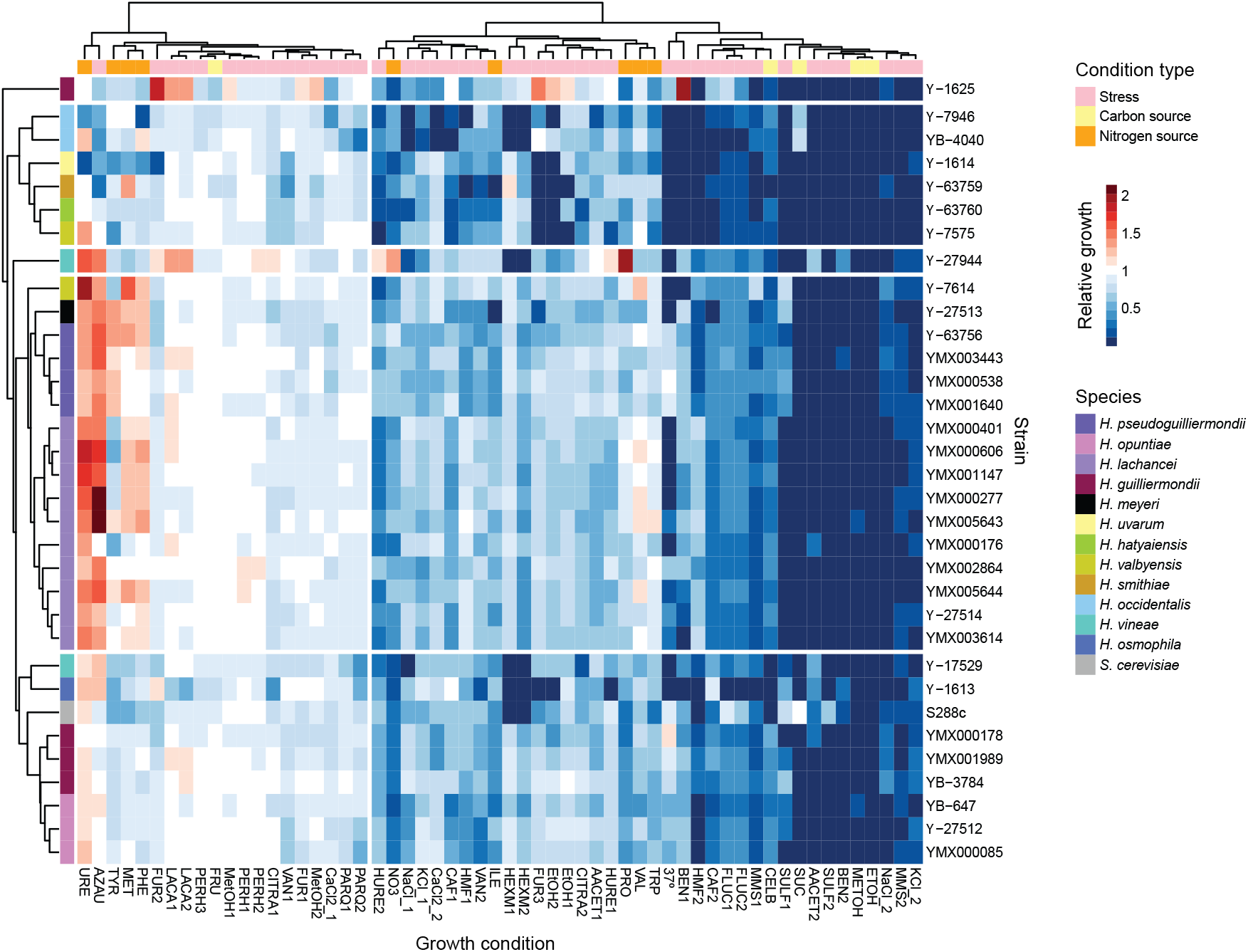
Phenotypic profiles of *Hanseniaspora* isolates. Heatmap showing hierarchical clustering of growth of the *Hanseniaspora* isolates from agave fermentations together with the reference strains. Growth was estimated as the ratio of the growth rate under each condition to the growth rate under rich medium and is depicted in a red (fast growth) to blue (slow growth) gradient as shown in the legend. Colored squares at the top of the heatmap show the type of condition, and squares at the left of the heatmap denote the species to which the strain belongs.

At the species level, hierarchical clustering revealed three distinct clusters. The first encompassed strains of *H. opuntiae, H. guilliermondii, H. osmophila* (Y-1613), one *H. vineae* strain (Y-17529), and *S. cerevisiae* S288C. The second cluster included strains of *H. lachancei, H. pseudoguilliermondii*, and one strain of *H. meyeri* (Y-27513) and one of *H. valbyensis* (Y-7614). Finally, the third cluster comprised strains of *H. occidentalis, H. uvarum, H. smithiae*, and one strain of *H. hatyaensis* (Y-63760) and one of *H. valbyensis* (Y-7575). Beyond these general patterns there were specific strains that displayed atypical performance. Notably, strains Y-27944 (*H. vineae*) and Y-1625 (*H. guilliermondii*) showed higher fitness under conditions which were generally detrimental to the other tested strains (e.g., presence of benomyl, furfural and lactic acid, Figure 5, Figure S3). Conversely, several strains exhibited markedly low fitness under several conditions. For example, strains Y-1613 (*H. osmophila*), Y-7946 (*H. occidentalis*) and YB-4040 (*H. occidentalis*).

We did not observe a clear association between the phenotypic profiles of the strains and the patterns of gene loss. Although individual strains exhibited variable growth responses across conditions, these differences did not correlate with the presence or absence of specific genes or functional categories. For example, the *H. lachancei* strains that have lost many genes associated with carbohydrate metabolism did not exhibit reduced fitness on the carbon sources that we tested. Similarly, the *H. opuntiae* strain from agave fermentations that is depleted in stress response genes is not particularly impaired to growth under conditions that impose cellular stresses. Therefore, our results do not support a direct role for gene loss in driving phenotypic diversity among *Hanseniaspora* isolates, suggesting that other mechanisms may underlie the observed phenotypic patterns.

## DISCUSSION

*Hanseniaspora* yeasts are an important group of organisms for the production of fermented beverages, including wine. In recent years, they have also attracted attention due to their accelerated rates of genomic change, particularly extensive gene losses. To gain deeper insight into the biology of this genus, we examined the interlineage genomic and phenotypic diversity within a collection of *Hanseniaspora* isolates obtained from traditional agave distilleries across Mexico. Previous work has shown that these yeasts are prevalent in agave fermentations (Gallegos-Casillas et al., 2024; Jara-Servin et al., 2025), and genome sequencing of the 15 strains reported here further revealed the coexistence of members of the quartet-species complex (*H. lachancei* - *H. pseudoguilliermondii* - *H. guilliermondii* - *H. opuntiae*) within this environment.

Comparative genomic analysis of the sequenced isolates aligns with the original findings of Steenwyk et al. (2019) regarding the extensive gene losses in *Hanseniaspora* species. Furthermore, our analyses revealed interlineage variation in gene content, particularly among isolates of *H. lachancei* and *H. opuntiae*. GO enrichment analysis also showed variation in gene loss within functional categories, both within and between species. The observed diversity of gene losses suggests that *Hanseniaspora* species continue to evolve through gene loss. This observation aligns with recent population studies from *Hanseniaspora* strains from other environments, which revealed intraspecific hybrids, varying levels of ploidy, and segmental duplications, showing the highly dynamic nature of these species’ genomes (Saubin et al., 2020; Onetto et al., 2025).

Phenotypic characterization of the *Hanseniaspora* isolates from agave fermentations revealed variation in how isolates respond to different carbon sources and stress conditions. However, these phenotypic patterns did not clearly correlate with the observed disparities in gene losses. In the only GWAS study performed in *Hanseniaspora* species, analysis of several *H. uvarum* isolates from wine revealed an association of coper tolerance with a single nucleotide mutation (Onetto et al., 2025). Whether selective pressures have contributed to the distinctive genomic features of *Hanseniaspora* species, particularly the loss of multiple genes related to cell cycle and genome integrity, remains an open question. One plausible explanation involves selection for rapid growth rates. As demonstrated in *H. uvarum*, fast growth is a key factor underlying its dominance during spontaneous grape juice fermentation (Onetto et al., 2025). Rapid proliferation could be facilitated by the elimination of genes associated with cell cycle regulation. Alternatively, the mutator strain hypothesis may account for the absence of genes involved in genome stability within *Hanseniaspora*. This model proposes that mutator strains become advantageous in scenarios of niche colonization or adaptation to variable environments, scenarios likely relevant to survival in agave fermentations (reviewed in Martínez-Cano et al., 2015; Oliver et al., 2000). Consequently, *Hanseniaspora* species may rely on a “runaway” evolutionary strategy that combines accelerated growth with elevated mutational rate. Noteworthy, this strategy bears a striking resemblance to the evolutionary dynamics observed in cancer cells (Natali and Rancati, 2019).

Given the prevalence of *Hanseniaspora* yeasts in open agave fermentations in Mexico, the populations in this region represent an exceptional model for investigating the evolutionary processes driving genome reduction and adaptation. By integrating genomic, ecological, and phenotypic analyses, future work can elucidate how these species persist under challenging environmental conditions despite extensive gene loss, providing broader insights into eukaryotic genome evolution in natural environments.

## DATA AVAILABILITY

Raw sequence reads have been deposited in the NCBI Sequence Read Archive under BioProject accession number PRJNA1182377.

## ACKNOWLEDGEMENTS

We are in debt to all the agave-spirit producers who welcomed us in their distilleries. More comprehensive acknowledgments of the producers can be found in Gallegos□Casillas, et. al. (2024). We thank Luis Aguilar and Santiago García for support and administration of computing resources, and Porfirio Gallegos□Casillas, Susana Ruiz-Castro, Carina Uribe and Alejandra Castillo for technical assistance.

## FUNDING

This work was funded by the Secretaría de Ciencia, Humanidades, Tecnología e Innovación del Gobierno de México (SECIHTI grants FORDECYT-PRONACES/103000/2020, CF-2023-G-695 and CBF-2025-G-838). This work was partially supported by UNAM-PAPIIT IN212524. LFG-O and EM were funded by SECIHTI at postdoctoral level (4133922) and for a sabbatical stay (I0200/111/2024), respectively. AMG-O was funded by the UK BBSRC under the Global Challenges Research Fund (GCRF) Growing Research Capability call through the CABANA Innovation Fund (BB/P027849/1).

## CONFLICT OF INTEREST

The authors declare that the research was conducted in the absence of any commercial or financial relationships that could be construed as a potential conflict of interest.

## MANUSCRIPT VERSIONS

This manuscript was released as a pre-print at *bioRxiv* (García-Ortega L. F., et al., 2025).

## SUPPLEMENTARY MATERIAL

Table S1. Summary of *Hanseniaspora* strains analyzed from agave fermentations.

Table S2. Reference genomes used for phylogenomic analysis.

Table S3. Strains assessed in the phenotyping assay.

Table S4. Culture conditions employed in the phenotyping assay.

Table S5. Assembly statistics of the *Hanseniaspora* genomes analyzed.

## Notes

### Competing Interest Statement

The authors have declared no competing interest.

