## Supplementary Material for "*Hanseniaspora* yeasts from agave fermentations provide insight into intraspecific gene loss"

Table S1. Summary of *Hanseniaspora* strains analyzed from agave fermentations.

| ID | Species MALDI | Score MALDI | ITS Species | ITS Identity | ITS Coverage | Species genome |
|---|---|---|---|---|---|---|
| YMX000538 | *H. opuntiae* | 2.17 | | | | *H. pseudoguilliermondii* |
| YMX003614 | *H. lachancei* | 2.08 | *H. lachancei* | 99.8 | 595/595 | *H. lachancei* |
| YMX003443 | *H. opuntiae* | 2.07 | *H. opuntiae* | 98.6 | 147/147 | *H. pseudoguilliermondii* |
| YMX000085 | *H. opuntiae* | 2.09 | *H. lachancei* | 98.9 | 641/641 | *H. opuntiae* |
| YMX000277 | *H. lachancei* | 1.97 | *H. lachancei* | 100.0 | 590/593 | *H. lachancei* |
| YMX000401 | *H. opuntiae* | 1.74 | *H. lachancei* | 99.8 | 642/642 | *H. lachancei* |
| YMX002864 | *H. lachancei* | 2.02 | *H. lachancei* | 99.8 | 618/618 | *H. lachancei* |
| YMX000176 | *H. lachancei* | 1.92 | *H. lachancei* | 100.0 | 593/594 | *H. lachancei* |
| YMX000178 | *H. opuntiae* | 1.51 | *H. guilliermondii* | 99.2 | 646/646 | *H. guilliermondii* |
| YMX001147 | *H. lachancei* | 2.17 | *H. lachancei* | 100.0 | 595/595 | *H. lachancei* |
| YMX001640 | *H. lachancei* | 1.92 | *H. opuntiae* | 98.9 | 646/650 | *H. pseudoguilliermondii* |
| YMX000606 | *H. lachancei* | 2.19 | *H. lachancei* | 99.8 | 639/639 | *H. lachancei* |
| YMX001989 | *H. opuntiae* | 1.60 | *H. guilliermondii* | 99.3 | 642/644 | *H. guilliermondii* |
| YMX005701 | *H. vineae* | 1.59 | *H. osmophila* | 99.3 | 158/159 | |
| YMX005697 | *H. opuntiae* | 1.93 | Not reliable | 97.7 | 268/269 | |
| YMX005644 | | | | | | *H. lachancei* |
| YMX005643 | | | | | | *H. lachancei* |

Table S2. Reference genomes used for phylogenomic analysis.

| Species | Strain | Genome version |
|---|---|---|
| *Hanseniaspora gamundiae* | CRUB 1928 | GCA_003020785.1 |
| *Hanseniaspora occidentalis* | CBS 6783 | GCA_030573535.1 |

| | | |
|---|---|---|
| *Hanseniaspora vineae* | T02/19AF | GCA_000585475.3 |
| *Hanseniaspora osmophila* * | AWRI3579 | GCA_001747045.1 |
| *Hanseniaspora singularis* | NRRL Y-63757 | GCA_030565715.1 |
| *Hanseniaspora smithiae* | CRUB 1602 | GCA_030573575.1 |
| *Hanseniaspora hatyaiensis* | ZIM 2327 | GCA_030573555.1 |
| *Hanseniaspora nectarophila* | CBS 13383 | GCA_030573495.1 |
| *Hanseniaspora thailandica* | ZIM 2325 | GCA_030573475.1 |
| *Hanseniaspora uvarum* | HU-10 | GCA_037102615.1 |
| *Hanseniaspora meyeri* * | APC 12.1 | GCA_030370665.1 |
| *Hanseniaspora clermontiae* | NRRL Y-27515 | GCA_030567235.1 |
| *Hanseniaspora guilliermondii* | NRRL Y-1625 | GCA_004919775.1 |
| *Hanseniaspora opuntiae* * | AWRI 3578 | GCA_001749795.1 |
| *Hanseniaspora pseudoguilliermondii* | ZIM213 | GCA_003708335.1 |
| *Hanseniaspora lachancei* | NRRL Y-27514 | GCA_004919765.1 |

* Genomes used for the construction of genetic models based on sequence homology.

Table S3. Strain assessed in the phenotyping assay.

| Species | ID | Substrate | Country |
|---|---|---|---|
| *H. opuntiae* | YMX000085 | Agave | Mexico |
| *H. lachancei* | YMX000176 | Agave | Mexico |
| *H. lachancei* | YMX000277 | Agave | Mexico |
| *H. lachancei* | YMX000401 | Agave | Mexico |
| *H. lachancei* | YMX000606 | Agave | Mexico |
| *H. lachancei* | YMX001147 | Agave | Mexico |
| *H. lachancei* | YMX002864 | Agave | Mexico |
| *H. lachancei* | YMX003614 | Agave | Mexico |
| *H. lachancei* | YMX005643 | Agave | Mexico |

| | | | |
|---|---|---|---|
| *H. lachancei* | YMX005644 | Agave | Mexico |
| *H. pseudoguilliermondii* | YMX003443 | Agave | Mexico |
| *H. pseudoguilliermondii* | YMX000538 | Agave | Mexico |
| *H. pseudoguilliermondii* | YMX001640 | Agave | Mexico |
| *H. guilliermondii* | YMX000178 | Agave | Mexico |
| *H. guilliermondii* | YMX001989 | Agave | Mexico |
| *H. opuntiae* | NRRL Y-27512$^{T}$ | Cactus rot | Hawaii Island |
| *H. opuntiae* | NRRL YB-647 | Tangerine | |
| *H. lachancei* | NRRL Y-27514$^{T}$ | Fermenting agave juice | Mexico |
| *H. pseudoguilliermondii* | NRRL Y-63756$^{T}$ | Orange juice concentrate | USA |
| *H. guilliermondii* | NRRL Y-1625$^{T}$ | Infected nail | South Africa |
| *H. guilliermondii* | NRRL YB-3784 | Yellow dates | USA |
| *H. valbyensis* | NRRL Y-7575 | Pulque | Mexico |
| *H. valbyensis* | NRRL Y-7614 | Canker, oak | USA |
| *H. uvarum* | NRRL Y-1614$^{T}$ | Muscatel grapes | Ukraine |
| *H. smithiae* | NRRL Y-63759$^{T}$ | *Cyttaria* spp. | Argentina |
| *H. hatyaiensis* | NRRL Y-63760$^{T}$ | Rotten wood | Thailand |
| *H. meyeri* | NRRL Y-27513$^{T}$ | Fruit of soapberry, *Sapindus* sp. | Hawaii Island |
| *H. vineae* | NRRL Y-17529$^{T}$ | Vineyard soil | South Africa |
| *H. vineae* | NRRL Y-27944 | Gut of dobsonfly, *Corydalus cornutus* | USA |
| *H. osmophila* | NRRL Y-1613$^{T}$ | Riesling grapes | Germany |
| *H. occidentalis* var. *occidentalis* | NRRL Y-7946$^{T}$ | Soil | West Indies |

| | | | |
|---|---|---|---|
| *H. occidentalis* var. *occidentalis* | NRRL YB-4040 | Tropical soil | Panama |

[T] Type strain

Table S4. Culture conditions employed in the phenotyping assay.

| Compound | Dose | Condition |
|---|---|---|
| | YPD (control medium) | |
| Furfural | 12.5 mM | FUR1 |
| Furfural | 25 mM | FUR2 |
| Furfural | 50 mM | FUR3 |
| Hydrogen peroxide | 0.5 mM | PERH1 |
| Hydrogen peroxide | 1.5 mM | PERH2 |
| Hydrogen peroxide | 3 mM | PERH3 |
| Methanol | 2 % | MetOH1 |
| Methanol | 5 % | MetOH2 |
| Ethanol | 6 % | EtOH1 |
| Ethanol | 12 % | EtOH2 |
| Acetic acid | 2 % | AACET1 |
| Acetic acid | 4 % | AACET2 |
| Citric acid | 20 mg/mL | CITRA1 |
| Citric acid | 40 mg/mL | CITRA2 |
| Lactic acid | 3 g/L | LACA1 |
| Lactic acid | 6 g/L | LACA2 |
| Menadione | 0.25 mM | MEN1 |
| Menadione | 0.5 mM | MEN2 |
| Cycloheximide | 0.01 % | HEXM1 |
| Cycloheximide | 0.1 % | HEXM2 |
| Caffeine | 2 mg/mL | CAF1 |

| | | |
|---|---|---|
| Caffeine | 4 mg/mL | CAF2 |
| Paraquat | 2.5 mM | PARQ1 |
| Paraquat | 5 mM | PARQ2 |
| Vainillin | 3 mM | VAN1 |
| Vainillin | 6 mM | VAN2 |
| Hydroxyurea | 5 mg/mL | HURE1 |
| Hydroxyurea | 15 mg/mL | HURE2 |
| Canavanine | 5 mg/mL | CANV1 |
| Benomyl | 50 mg/L | BEN1 |
| Benomyl | 100 mg/L | BEN2 |
| Fluconazol | 60 mg/mL | FLUC1 |
| Fluconazol | 150 mg/mL | FLUC2 |
| Hygromycin b | 100 mg/mL | HYGRO1 |
| Hygromycin b | 200 mg/mL | HYGRO2 |
| Methyl methanesulfonate | 0.02 % | MMS1 |
| Methyl methanesulfonate | 0.04 % | MMS2 |
| KCl | 1 M | KCl_1 |
| KCl | 2 M | KCl_2 |
| LiCl | 125 mM | LiCl_1 |
| LiCl | 250 mM | LiCl_2 |
| NaCl | 0.75 M | NaCl_1 |
| NaCl | 1.5 M | NaCl_2 |
| $CaCl_2$ | 100 mM | $CaCl_2$_1 |
| $CaCl_2$ | 200 mM | $CaCl_2$_2 |
| $Na_2SO_3$ | 40 mM | SULF1 |
| $Na_2SO_3$ | 80 mM | SULF2 |
| Heat | 37 ºC | G_37 |

| | | |
|---|---|---|
| YPD_DMSO (0.75 %) (control medium) | | |
| Hydroxymethylfurfural | 12.5 mM | HMF1 |
| Hydroxymethylfurfural | 25 mM | HMF2 |
| YNB-Glucose (2 %) (control medium) | | |
| Galactose | 2 % | GAL |
| Sucrose | 2 % | SUC |
| Maltose | 2 % | MAL |
| Cellobiose | 2 % | CELB |
| Melibiose | 2 % | MELI |
| Fructose | 2 % | FRU |
| Glycerol | 2 % | GLYC |
| Sorbitol | 2 % | SORB |
| Raffinose | 2 % | RAF |
| Lactose | 2 % | LAC |
| Xylose | 2 % | XYL |
| Methanol | 2 % | METOH |
| Ethanol | 2 % | ETOH |
| Acetate | 2 % | ACET |
| Citrate | 2 % | CITR |
| YNB_$NH_4$ (Glucose 2 %) (control medium) | | |
| Valine | 0,01 g/L (N) | VAL |
| Methionine | 0,01 g/L (N) | MET |
| Tyrosine | 0,01 g/L (N) | TYR |
| Isoleucine | 0,01 g/L (N) | ILE |
| Leucine | 0,01 g/L (N) | LEU |
| Phenylalanine | 0,01 g/L (N) | PHE |
| Tryptophan | 0,01 g/L (N) | TRP |

| | | |
|---|---|---|
| Proline | 0,01 g/L (N) | PRO |
| Urea | 0,01 g/L (N) | URE |
| Sodium nitrite | 0,01 g/L (N) | NO2 |
| Sodium nitrate | 0,01 g/L (N) | NO3 |
| 6-Azauracil | 500 mg/mL | AZAU |

Table S5. Assembly statistics of the *Hanseniaspora* genomes analyzed.

| Specie/Strain | Number of contigs | Number of scaffolds | Total size (bp) | Average GC content (%) | Number of genes | Average gene length | Number of exons | BUSCO completeness (genome) | BUSCO completeness (annotation) | Repetitive sequences (bp) |
|---|---|---|---|---|---|---|---|---|---|---|
| *H. lachancei* NRRL Y-27514 | 3,155 | 432 | 8,900,485 | 31.55 | 4,351 | 1,476 | 4,624 | 56.9 | 80.2 | 186,639 (2.10%) |
| YMX 004426 | 3,999 | 56 | 8,760,495 | 35.10 | 3,823 | 1,496 | 4,155 | 59.0 | 76.1 | 148,672 (1.70%) |
| YMX000176 | 1,272 | 68 | 8,784,353 | 35.07 | 4,177 | 1,510 | 4,389 | 59.1 | 81.1 | 164,723 (1.88%) |
| YMX000277 | 1,003 | 70 | 8,775,666 | 35.08 | 4,168 | 1,513 | 4,411 | 59.3 | 81.1 | 143,783 (1.64%) |
| YMX000401 | 1,937 | 82 | 8,791,455 | 35.09 | 4,173 | 1,514 | 4,395 | 59.0 | 81.1 | 144,215 (1.64%) |
| YMX000606 | 1,169 | 64 | 8,772,893 | 35.09 | 4,168 | 1,513 | 4,399 | 59.1 | 81.1 | 143,577 (1.64%) |
| YMX001147 | 1,153 | 73 | 8,778,582 | 35.09 | 4,157 | 1,521 | 4,383 | 58.6 | 81.4 | 144,343 (1.64%) |
| YMX005643 | 800 | 31 | 8,755,234 | 35.11 | 3,824 | 1,519 | 3,356 | 59.2 | 76.2 | 144,049 (1.65%) |
| YMX005644 | 2,018 | 62 | 8,746,122 | 35.11 | 3,829 | 1,497 | 4,176 | 59.8 | 76.2 | 147,677 (1.69%) |
| YMX003614 | 2,127 | 97 | 8,778,599 | 35.07 | 4,173 | 1,513 | 4,434 | 58.6 | 81.0 | 144,628 (1.65%) |
| YMX002864 | 2,039 | 85 | 8,835,134 | 35.05 | 4,175 | 1,524 | 4,397 | 59.4 | 81.6 | 153,083 (1.73%) |
| *H. pseudoguilliermondii* ZIM213 | 917 | 917 | 9,077,954 | 30.91 | 4,297 | 1,506 | 4,590 | 58.5 | 80.9 | 436,146 (4.80%) |
| YMX000538 | 2,693 | 282 | 8,711,511 | 34.46 | 4,235 | 1,489 | 4,422 | 58.7 | 80.3 | 149,358 (1.71%) |
| YMX001640 | 2,782 | 212 | 8,799,227 | 34.41 | 4,251 | 1,496 | 4,470 | 59.8 | 80.8 | 167,394 (1.90%) |
| YMX003443 | 2,513 | 233 | 8,809,252 | 34.42 | 4,246 | 1,496 | 4,464 | 58.8 | 80.7 | 168,853 |

| | | | | | | | | | | |
|---|---|---|---|---|---|---|---|---|---|---|
| | | | | | | | | | | (1.92%) |
| *H. opuntiae* AWRI3578 | 17 | 17 | 8,831,957 | 31.56 | 4,244 | 1,533 | 4,524 | 56.9 | 80.3 | 162,890 (1.84%) |
| YMX000085 | 5,725 | 1,116 | 8,854,817 | 34.83 | 4,492 | 1,344 | 4,689 | 51.8 | 72.1 | 169,016 (1.91%) |
| *H. guilliermondii* NRRL Y-1625 | 395 | 395 | 9,135,640 | 27.24 | 4,195 | 1,529 | 4,484 | 59.5 | 81.3 | 349,695 (3.83%) |
| YMX000178 | 4,202 | 31 | 9,069,783 | 30.98 | 4,187 | 1,526 | 4,437 | 61.3 | 81.8 | 306,488 (3.38%) |
| YMX001989 | 883 | 28 | 9,062,411 | 30.98 | 4,176 | 1,530 | 4,424 | 60.9 | 81.9 | 292,076 (3.22%) |
